# Structural Brain Imaging Studies Offer Clues about the Effects of the Shared Genetic Etiology among Neuropsychiatric Disorders

**DOI:** 10.1101/809582

**Authors:** Nevena V. Radonjić, Jonathan L. Hess, Paula Rovira, Ole Andreassen, Jan K. Buitelaar, Christopher R. K. Ching, Barbara Franke, Martine Hoogman, Neda Jahanshad, Carrie McDonald, Lianne Schmaal, Sanjay M. Sisodiya, Dan J. Stein, Odile A. van den Heuvel, Theo G.M. van Erp, Daan van Rooij, Dick J. Veltman, Paul Thompson, Stephen V. Faraone

## Abstract

**Background:** Genomewide association studies have found significant genetic correlations among many neuropsychiatric disorders. In contrast, we know much less about the degree to which structural brain alterations are similar among disorders and, if so, the degree to which such similarities have a genetic etiology.

**Methods:** From the Enhancing Neuroimaging Genetics through Meta-Analysis (ENIGMA) consortium, we acquired standardized mean differences (SMDs) in regional brain volume and cortical thickness between cases and controls. We had data on 41 brain regions for: attention-deficit/hyperactivity disorder (ADHD), autism spectrum disorder (ASD), bipolar disorder (BD), epilepsy, major depressive disorder (MDD), obsessive compulsive disorder (OCD) and schizophrenia (SCZ). These data had been derived from 24,360 patients and 37,425 controls.

**Results:** The SMDs were significantly correlated between SCZ and BD, OCD, MDD, and ASD. MDD was positively correlated with BD and OCD. BD was positively correlated with OCD and negatively correlated with ADHD. These pairwise correlations among disorders were correlated with the corresponding pairwise correlations among disorders derived from genomewide association studies (*r* = 0.49).

**Conclusions:** Our results show substantial similarities in sMRI phenotypes among neuropsychiatric disorders and suggest that these similarities are accounted for, in part, by corresponding similarities in common genetic variant architectures.

## Introduction

Neuropsychiatric disorders have substantial heritability, as shown by many studies of twins and families (1). Genomewide association studies (GWAS) have shown that common genetic variants account for some of this heritability, and that some of this heritability is shared across neuropsychiatric disorders (2-5). The genetic overlap across disorders may partly explain why these disorders tend to co-occur with one another in both clinical and community samples (6).

Subcortical brain volumes and cortical thickness/surface area dynamically change from early development through adulthood and old age. A study of the Enhancing Neuroimaging Genetics through Meta-Analysis (ENIGMA) Plasticity Working Group reported that changes in structural magnetic resonance imaging (sMRI) phenotypes have heritabilities ranging from 5% for pallidum to 42% for cerebellar gray matter (7). Heritability estimates of change rates were age-related and generally higher in adults than in children, probably due to an increasing influence of genetic factors with age (7). ENIGMA sMRI studies of different psychiatric and neurological disorders further characterized MRI-derived phenotypes that can be used to assess heritability (reviewed in 8).

ENIGMA has also reported significant case vs. control differences in sMRI phenotypes for: attention-deficit/hyperactivity disorder (ADHD) (9, 10), autism spectrum disorder (ASD) (11), bipolar disorder (BD) (12, 13), common epilepsy syndromes (14), major depressive disorder (MDD) (15, 16), obsessive compulsive disorder (OCD) (17, 18) and schizophrenia (SCZ) (19, 20). Here we estimate the degree of similarity in sMRI phenotypes among these disorders and evaluate whether these similarities are influenced by corresponding similarities in common genetic variant architectures.

## Methods

### Collection of structural neuroimaging summary statistics

Summary statistics from ENIGMA structural neuroimaging studies were collected from 12 multi-site analyses published by the ENIGMA Consortium for the following neuropsychiatric disorders: ADHD (9, 10), ASD (11), BD (12, 13), epilepsy (14), MDD (15, 16), OCD (17, 18), and SCZ (19, 20). Prior to computing the summary statistics, the regional brain volumes had been segmented with a common ENIGMA protocol using FreeSurfer software. Each site performed these segmentations on their raw data. In addition, quality control protocols provided by ENIGMA were run at each site. Details are at: http://enigma.ini.usc.edu/protocols/imaging-protocols.

The ADHD and ASD samples comprised both youth and adults. The other samples comprised adults only. The ethnicity of the patients was not available for all participants. The “epilepsy” cohort comprised temporal lobe epilepsy, genetic generalized epilepsy, and extra temporal epilepsy. We analyzed 7 subcortical and 34 cortical regions (total of 41 brain regions; the mean of left and right structures) that were included in the above specified ENIGMA studies. We extracted the covariate-adjusted Cohen’s *d* standardized mean differences (SMDs) denoting the case versus unaffected comparison subject differences in subcortical volume and cortical thickness/surface area measures. The covariates used in these studies adjusted SMDs for several covariates as indicated in Supplemental Table 1.

**Table 1.**
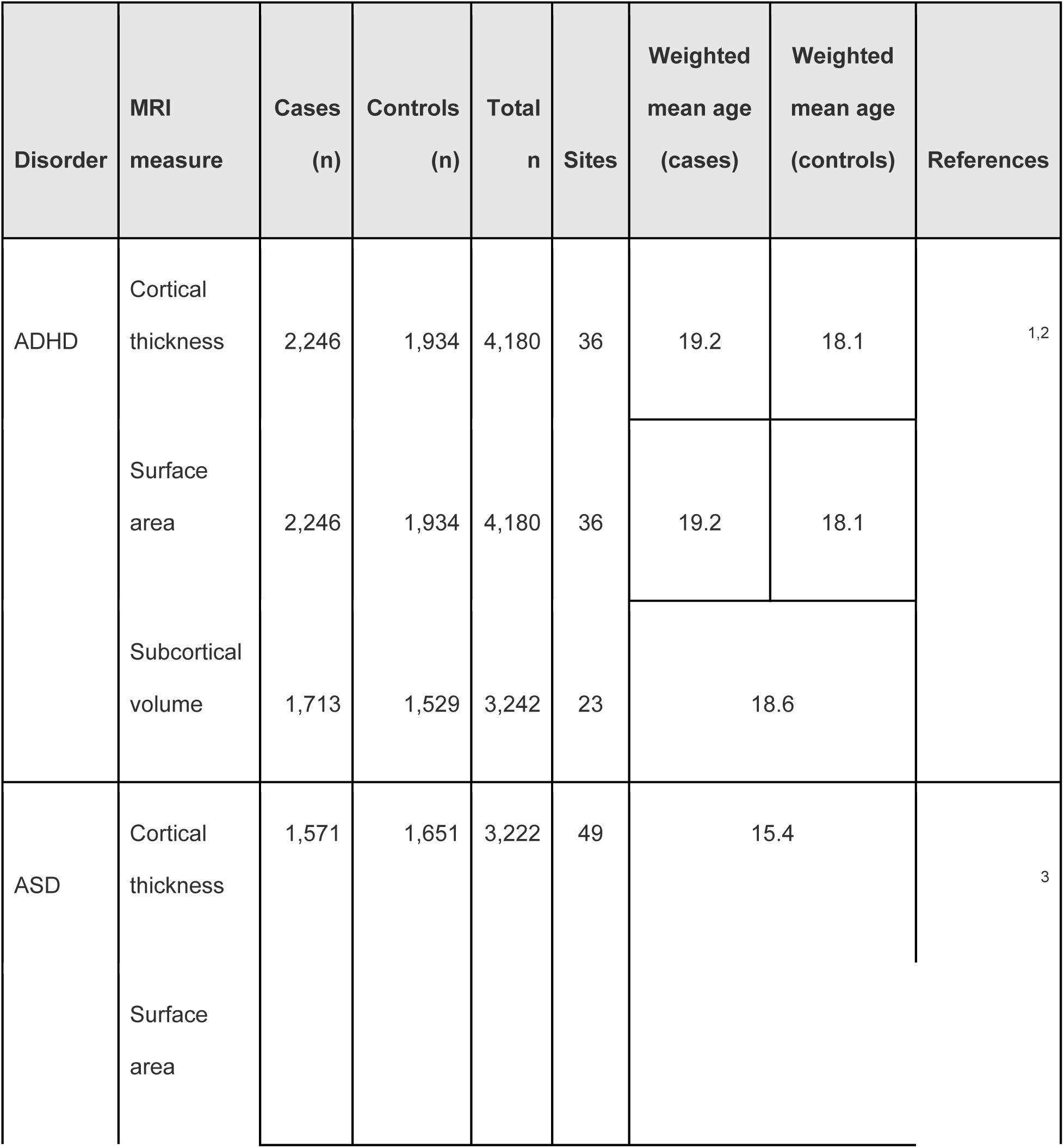
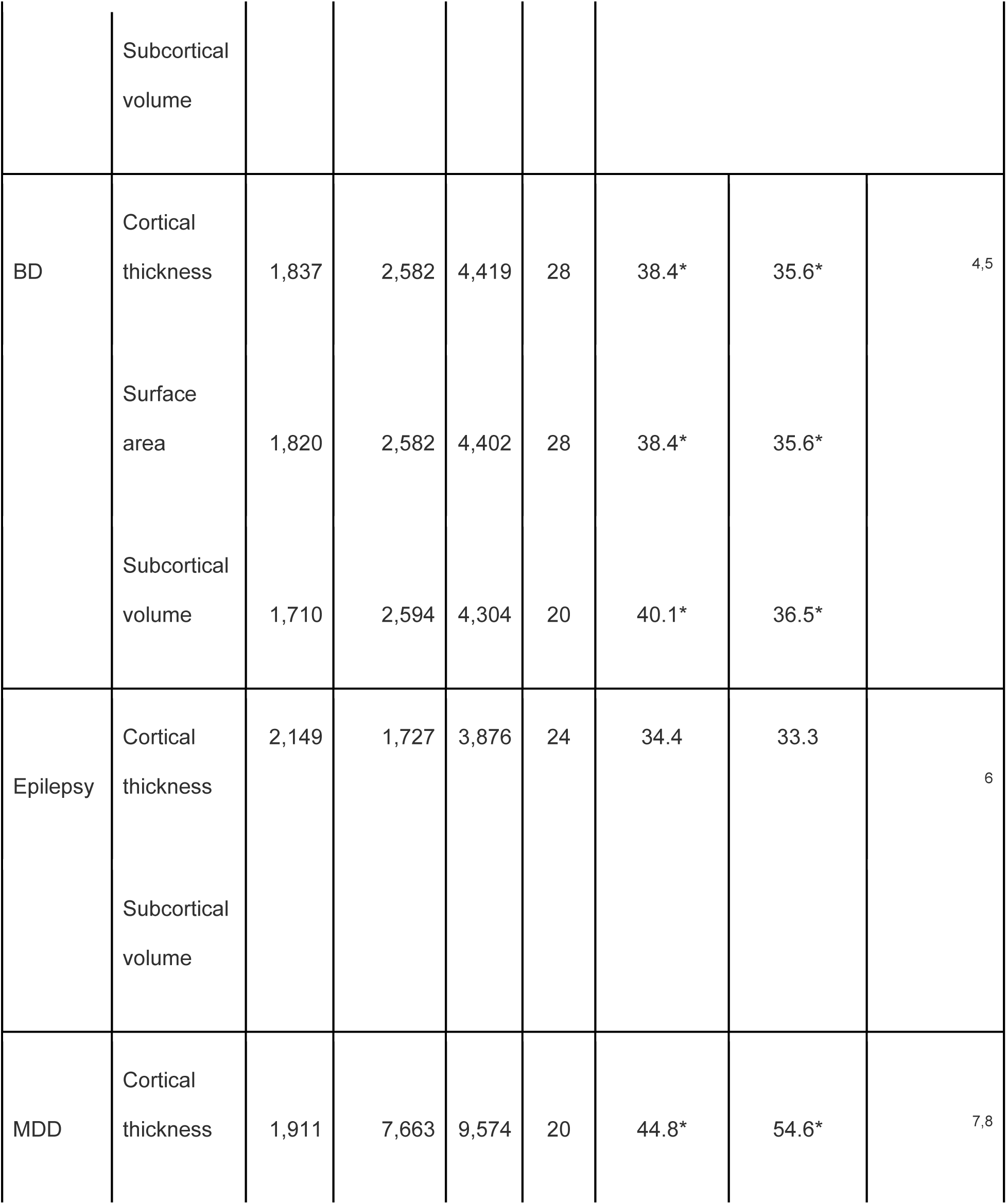
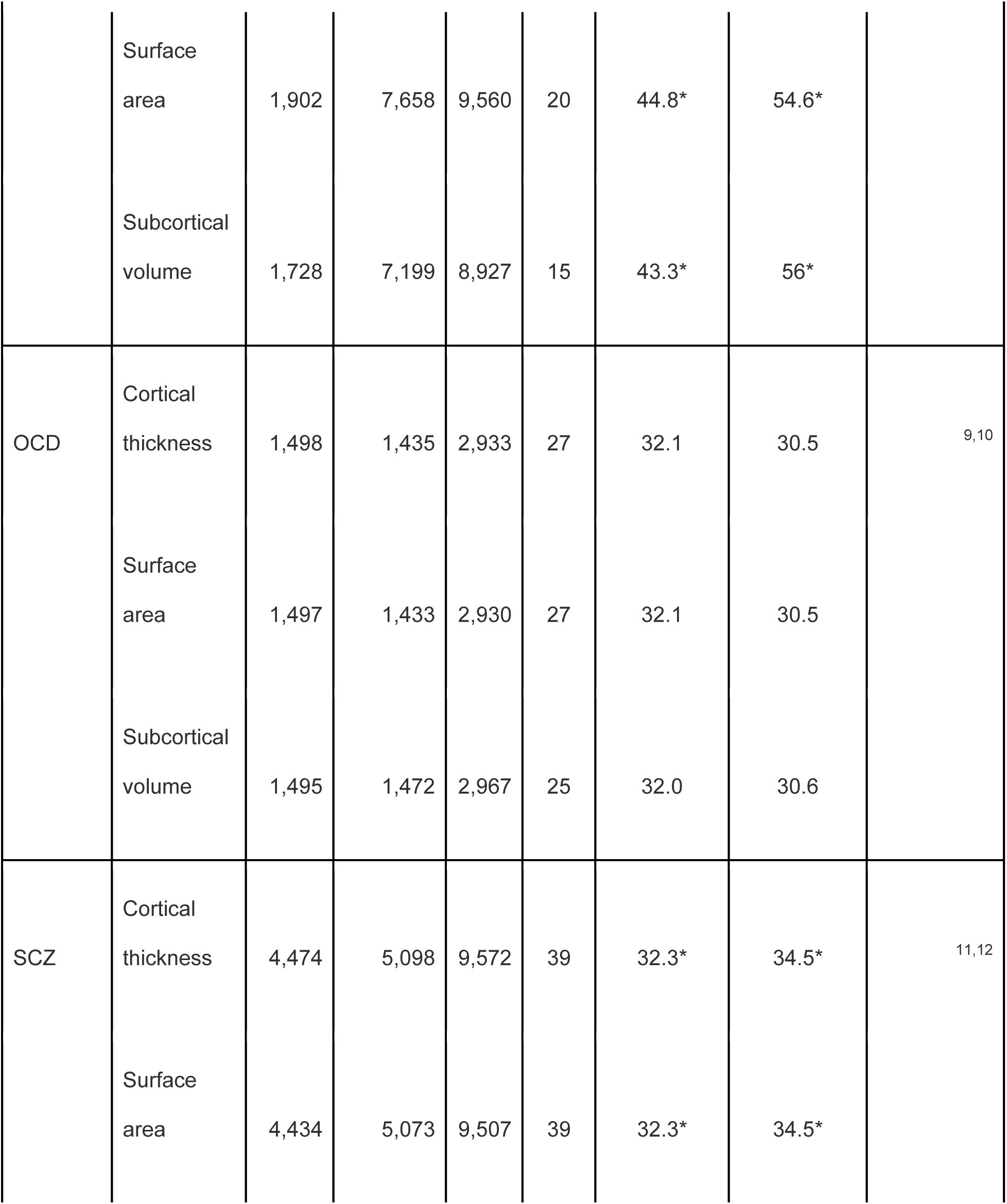
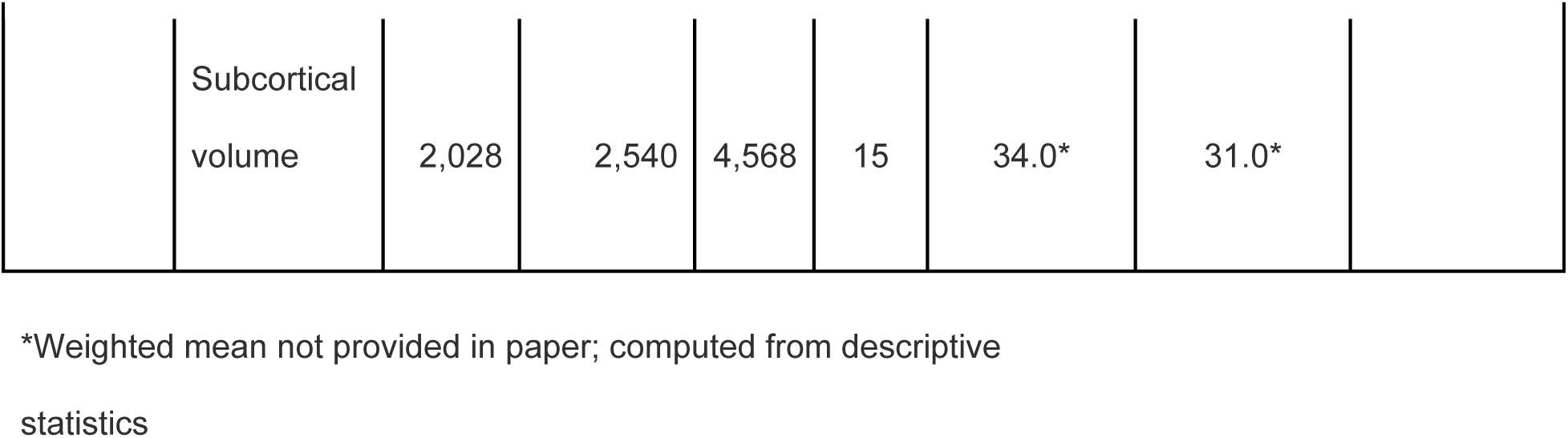
Sample demographics for the twelve studies by the ENIGMA Consortium into structural brain alterations in neuropsychiatric disorders.

#### Collection of GWAS results among neuropsychiatric disorders

Publicly available summary statistics from GWAS were downloaded from the Psychiatric Genomics Consortium (PCG) website (https://www.med.unc.edu/pgc/results-and-downloads/) with the exception of GWAS results for MDD coming from an online resource hosted by the University of Edinburgh (http://dx.doi.org/10.7488/ds/2458) and of GWAS results for epilepsy coming from the online Epilepsy Genetic Association Database (epiGAD) (http://www.epigad.org/gwas_ilae2018_16loci.html). Presented in Supplementary Table 2 are the numbers of affected cases and unaffected control participants included in each GWAS.

**Table 2.** Cross-disorder structural MRI phenotype correlations (ordered from smallest to largest *p*-value) based on Cohen’s d values obtained from the ENIGMA Project.

|  |  | sMRI correlation |  |  |  | Boferroni adjusted *p*- |
| --- | --- | --- | --- | --- | --- | --- |
| Disorder 1 | Disorder 2 | Pearson's *r* | df | se | *p*-value | value |
| BD | SCZ | 0.81 | 73 | 0.068 | 1.13E-18 | 2.38E-17 |
| BD | MDD | 0.69 | 73 | 0.085 | 1.21E-11 | 2.55E-10 |
| OCD | SCZ | 0.65 | 72 | 0.090 | 5.53E-10 | 1.16E-08 |
| MDD | SCZ | 0.58 | 73 | 0.095 | 5.55E-08 | 1.17E-06 |
| ADHD | BD | -0.53 | 73 | 0.099 | 1.18E-06 | 2.48E-05 |
| BD | OCD | 0.50 | 72 | 0.102 | 4.74E-06 | 9.95E-05 |
| MDD | OCD | 0.46 | 72 | 0.104 | 3.28E-05 | 6.89E-04 |
| ASD | BD | 0.38 | 73 | 0.108 | 8.98E-04 | 0.02 |
| ASD | SCZ | 0.36 | 73 | 0.109 | 1.35E-03 | 0.03 |
| ADHD | MDD | -0.33 | 73 | 0.111 | 4.27E-03 | 0.09 |
| ADHD | SCZ | -0.32 | 73 | 0.111 | 4.63E-03 | 0.10 |
| Epilepsy | MDD | -0.37 | 39 | 0.149 | 0.02 | 0.38 |
| ADHD | Epilepsy | -0.36 | 39 | 0.149 | 0.02 | 0.41 |
| ASD | MDD | 0.26 | 73 | 0.113 | 0.02 | 0.46 |
| Epilepsy | OCD | -0.19 | 39 | 0.157 | 0.23 | 1 |
| BD | Epilepsy | 0.17 | 39 | 0.158 | 0.30 | 1 |
| ADHD | OCD | -0.10 | 72 | 0.117 | 0.39 | 1 |
| ADHD | ASD | -0.06 | 73 | 0.117 | 0.60 | 1 |
| Epilepsy | SCZ | -0.03 | 39 | 0.160 | 0.86 | 1 |
| ASD | Epilepsy | 0.02 | 39 | 0.160 | 0.91 | 1 |
| ASD | OCD | 0.00 | 72 | 0.118 | 0.97 | 1 |

Note, the full meta-analysis GWAS of MDD that included data from 23andMe was not available for public release, thus we used the meta-analysis that combined results from the PGC cohorts and UK Biobank.

### Genetic and sMRI phenotype correlations among neuropsychiatric disorders

Linkage disequilibrium (LD)-score regression, a popular approach designed to analyze summary statistics from GWAS, was used to quantify the amount of shared genetic heritability, or genetic correlation (*r*_g_), existing between pairs of neuropsychiatric disorders, considering HapMap3 LD-scores (21).For these analyses, the largest and latest GWAS available for each neuropsychiatric disorder was selected and filtered to exclude markers with INFO<0.90 or within the MHC region (hg19:chr6:25-35Mb) (Supplementary Table 1).

To derive an estimate of the degree to which sMRI phenotypes were similar among disorders, we computed pairwise Pearson’s correlation coefficients between the Cohen’s d SMDs for each pair of disorders. We then used Pearson’s correlation to estimate, whether the genetic correlations for each disorder covaried with the sMRI phenotype correlations. However, a reliable *p*-value could not be calculated due to the lack of statistical independence among these pairs of correlation. Adjustments for sample overlap would have been possible with individual-level data, but the present study only had access to summary statistics. In a leave-one-out analysis, we iteratively excluded one pair of disorder correlations from the set and recalculated Pearson’s correlation coefficients to determine whether correlations were driven by any pair of disorders. We used classical multidimensional scaling (MDS) with correlation as the distance measure to visualize and help interpret the sMRI phenotype correlations. MDS summarizes the correlations among disorders in their SMDs by plotting them in a low-dimensional space for which the distance between disorders is proportional to their correlations.

Binomial sign tests were used to determine whether the number of disorders showing the same direction of effect in the sMRI phenotypes was greater than expected by chance (null probability of 50%). Per brain region, we performed Cochran’s *Q* test implemented in the *R* package *metafor* (v.2.1-0) to determine whether variability among Cohen’s d values was greater than expected by chance. All statistical analyses were performed with *R* version 3.5.2 (R Core Team, 2018), except for multidimensional scaling, for which we used STATA15 (22). We adjusted for repeated correlation tests using the Bonferroni procedure. Correlations showing a Bonferroni-adjusted p < 0.05 were considered significant (threshold *p* = 0.00227).

## Results

Sample demographics for the twelve studies by the ENIGMA Consortium on structural brain abnormalities in neuropsychiatric disorders are presented in Table 1.

### Case-control differences in subcortical volume and cortical surface area and thickness within neuropsychiatric disorders

Figure 1 presents a heatmap graph showing standardized effect sizes (Cohen’s d) measuring alterations in subcortical volume, cortical surface area and cortical thickness for 41 brain regions within seven neuropsychiatric disorders – ADHD, ASD, OCD, epilepsy, MDD, BD and SCZ. These have been reported on prior publications. The variation in color from blue to red illustrates the phenomenon of SBRV, with some regions showing significant reductions (blue) in volume/thickness/ surface areas and others not being affected. As indicated by the blueness of the cells, the most prominent reductions were seen for SCZ (mean Cohen’s d across all regions = −0.22, SE = 0.014), epilepsy (mean Cohen’s d = −0.12, SE = 0.017) and BD (mean Cohen’s d =-0.097, SE = 0.011). The smallest changes were observed for MDD (mean Cohen’s d = −0.018, SE = 0.006). All regions except for the caudate and putamen exhibited significant differences in the magnitude of Cohen’s d across disorders (Cochran’s Q p-values = 0.012 – 2.8×10^−32^). Eighteen sMRI phenotypes exhibited homogeneity with respect to sign of Cohen’s d across each of the neuropsychiatric disorders evaluated (binomial sign test *p*-values < 0.05): cortical thicknesses for caudal middle frontal gyrus, entorhinal cortex, fusiform gyrus, inferior temporal gyrus, insula, lateral orbitofrontal cortex, lingual gyrus, middle temporal gyrus, paracentral lobule, parahippocampal gyrus, pars opercularis of inferior temporal gyrus, precentral gyrus, precuneus, rostral anterior cingulate cortex, and supramarginal gyrus; subcortical volume for the hippocampus; and surface area for middle temporal gyrus, pars triangularis of inferior temporal gyrus, and pericalcarine cortex. For sMRI phenotypes for 39 regions of interest varying degrees of heterogeneity were noted in terms of discrepancy of signs of Cohen’s d. For example, individuals with ASD showed a slightly thicker cortex in the rostral middle frontal gyrus, individuals with ADHD showed no difference, and all other disorders showed a thinner cortex in this region compared to controls.

**Figure 1.**
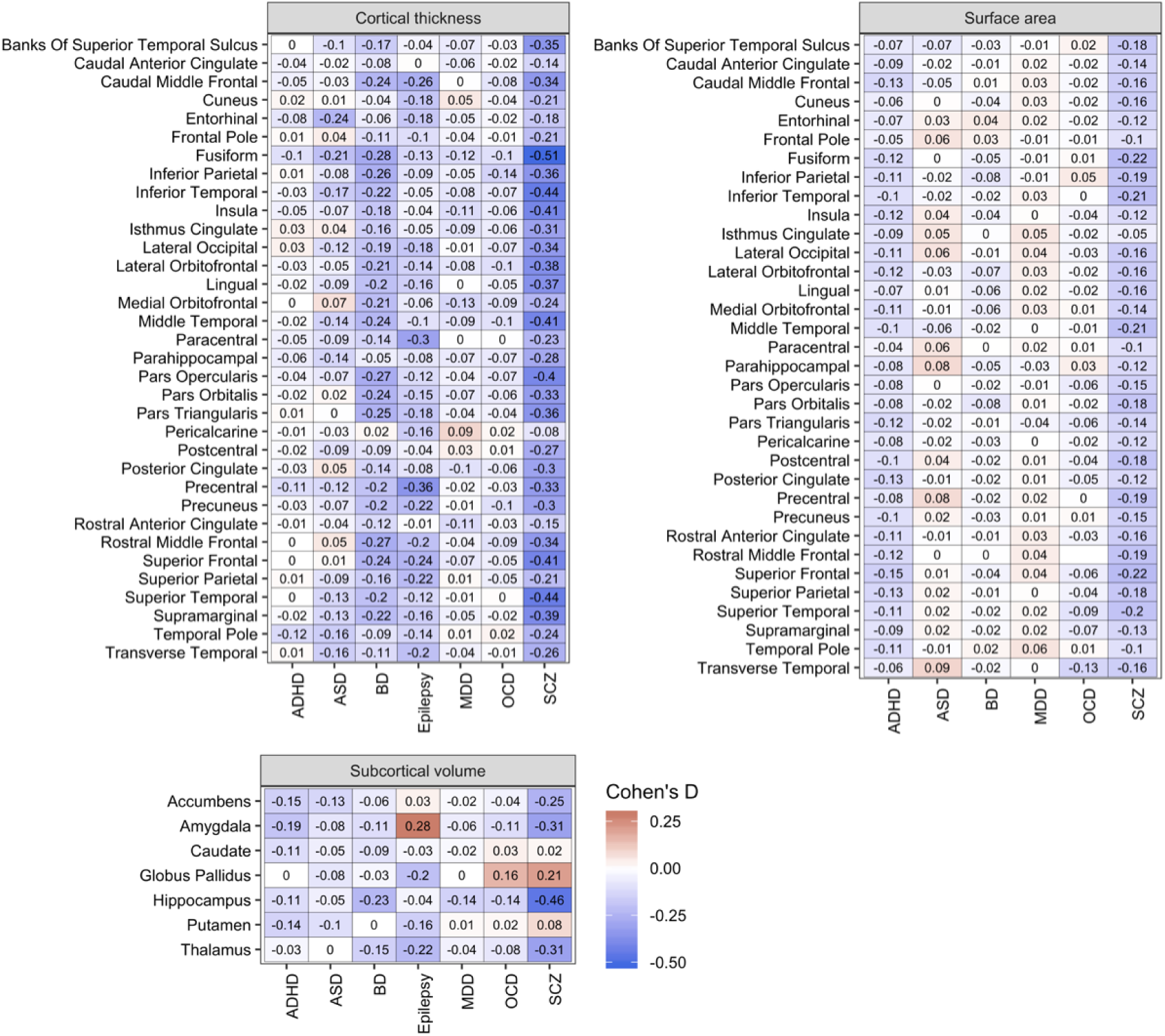
Case-control differences in subcortical volume and cortical thickness and surface area within neuropsychiatric disorders: A heatmap showing standardized mean differences (Cohen’s d) measuring case-control differences in subcortical volumes and cortical thickness for seven neuropsychiatric disorders. Results were obtained from ENIGMA working group publications. Negative values for Cohen’s d indicate smaller sizes of brain regions in cases versus unaffected comparisons. Note: ADHD – attention-deficit/hyperactivity disorder; ASD – autism spectrum disorder; BD – bipolar disorder; MDD – major depressive disorder; OCD – obsessive compulsive disorder; SCZ – schizophrenia.

### sMRI phenotype correlations among neuropsychiatric disorders

For each pair of disorders, we computed the Pearson correlation between their sMRI phenotypes listed in Figure 1. These are listed in Table 2, sorted by the magnitude of the correlation. The highest positive correlation was between SCZ and BD (*r*=0.81, *df* = 73, *p*<1.3×10^−18^, Bonferroni *p*=2.38×10^−17^). SCZ was also positively correlated with OCD (*r*=0.65, *df* = 72, *p*=5.5×10^−10^, Bonferroni *p*=1.2×10^−8^), ASD (*r* = 0.36, *df* = 73, *p* = 0.0014, Bonferroni *p* = 0.03), and MDD (*r*=0.57, *df* = 73, *p*=5.5×10^−8^, Bonferroni *p*=1.2×10^−6^). MDD was positively correlated with BD (*r*=0.68, *df* = 73, *p*=1.2×10^−11^, Bonferroni *p*=2.5×10^−10^) and OCD (*r*=0.46, *df* = 72, *p*=3.3×10^−5^, Bonferroni *p*=6.9×10^−4^). BD was positively correlated with OCD (*r*=0.50, *df* = 72, *p*=4.7×10^−6^, Bonferroni *p*=9.9×10^−5^) and ASD (*r*=0.38, *df* = 73, *p*=9.0×10^−4^, Bonferroni *p* = 0.02), and negatively correlated with ADHD (*r*=-0.53, *df* = 73, *p*=1.2×10^−6^, Bonferroni *p*=2.5×10^−5^). There were a few additional nominally significant negative correlations, which did not survive multiple testing correction: MDD and epilepsy (*r*=-0.37, *p*=0.02), MDD and ADHD (*r*=-0.33, *p*=0.004), SCZ and ADHD (*r*=-0.32, *p*=0.005), ADHD and epilepsy (*r*=-0.36, *p*=0.02), and a positive correlation between MDD and ASD (*r* = 0.26, *p* = 0.02).

Figure 2 visualizes the cross-disorder sMRI phenotype correlations by presenting the MDS configuration. We chose a three-dimensional solution, which accounted for 96.3% of the variation in the sMRI phenotype correlations. The Shepard diagram (Figure 2a) shows a good correspondence between the actual correlations and those predicted by the scaling solution. The three configuration plots illustrate the cross-disorder similarity in sMRI brain phenotypes (Figures 2b, c & d). Figure 2b shows the configuration for the first two dimensions, which jointly account for 86.6% of the variation in the data. It shows a clear clustering of ADHD, OCD, ASD and MDD that is distinct from epilepsy, SCZ and BD. The latter two disorder are similar on dimension 2 but differ on dimension 1. Figures 2c and 2d visualize the effect of adding the third dimension, which accounts for only 9.7% of the variance. This third dimension accounts for variance that makes the two mood disorders similar to one another and separates ADHD from other disorders.

**Figure 2:**
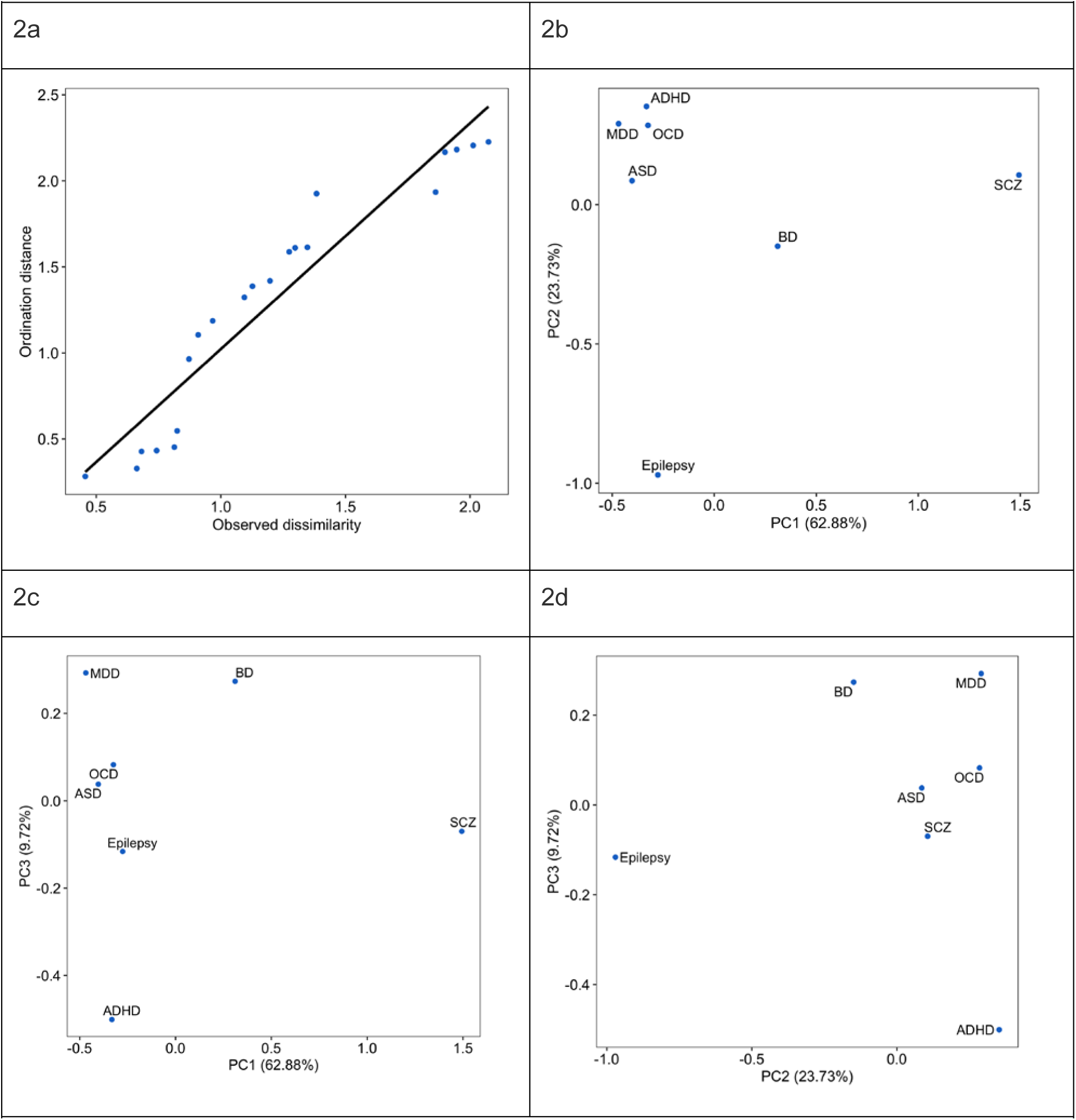
Multidimensional Scaling Configuration of sMRI Phenotype Cross Disorder Correlations: The Shepard diagram (Figure 2a) shows correspondence between the actual correlations and those predicted by the scaling solution. The three configuration plots illustrate the cross-disorder similarity in sMRI brain phenotypes according to principal component values (Figures 2b, c & d). Note: ADHD – attention-deficit/hyperactivity disorder; ASD – autism spectrum disorder; BD – bipolar disorder; MDD – major depressive disorder; OCD – obsessive compulsive disorder; SCZ – schizophrenia; PC = Principal Component.

### Correlation of shared genetic heritability with brain structural correlation

Figure 3 shows the pairwise correlations of sMRI phenotypes and genetic overlap across each pair of neuropsychiatric disorders. The vertical axis represents the between-disorder LD-score genetic correlations obtained from the PGC studies. The horizontal axis represents the between-disorder Cohen’s d value correlations for sMRI abnormalities obtained from the ENIGMA studies. Each dot represents genetic correlation and the Cohen’s d value correlation pairs for disorders as indicated by the legend (e.g., SCZ and BD, represented in the top right corner, show high genetic correlations and high correlations among their structural phenotype abnormalities compared to controls). The LD-score cross-disorder genetic correlations are positively correlated with the sMRI phenotype cross-disorder correlations (*r* = 0.49), thus we approximated that 24% of variance (measured by *R*^2^) in cross-disorder sMRI similarity can be accounted for by genetic correlations. Leave-one-out sensitivity analyses confirmed that the direction of the correlation remained positive and roughly at the same magnitude despite removal of individual pairs of disorders from the correlation test (range of Pearson’s *r* = 0.35 – 0.60), except for removing SCZ/BD (Pearson’s *r* = 0.35). SCZ and BD showed the highest degree of concordance with respect to genetic and sMRI phenotype correlations. We did a second sensitivity analysis that removed one disorder at a time. The results show an increase in the correlation when ADHD is removed and a decrease when schizophrenia is removed (Supplemental Figure 1).

**Figure 3.**
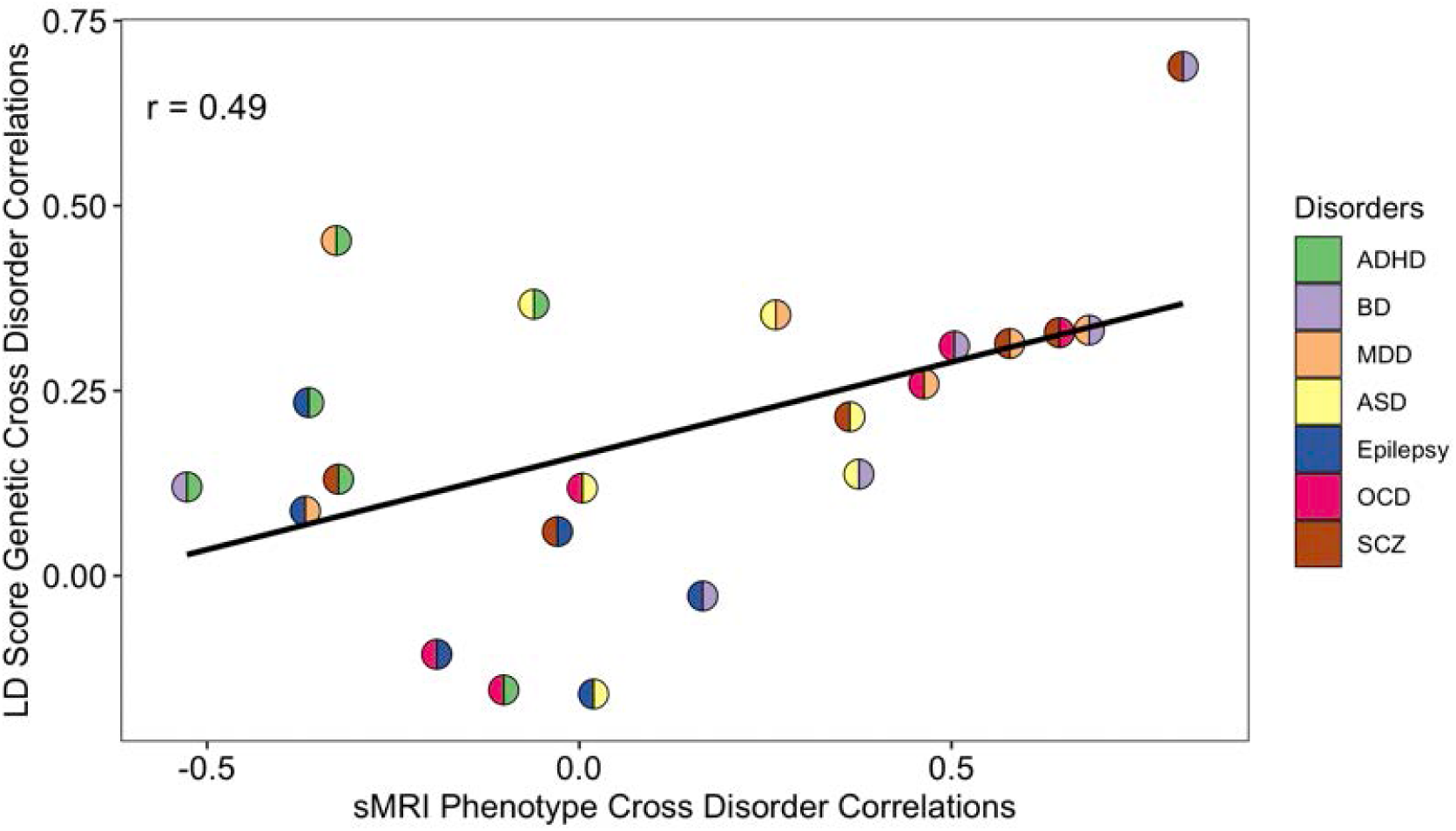
Correlation of shared genetic heritability with brain structural correlation: Scatter plot showing the correlation of correlations. Genetic correlations (*r*_g_) computed by LD-score regression are on the vertical axis, with correlations of Cohen’s d values displayed on the horizontal axis. Each dot is color-coded according to the pair-wise disorder correlations that were computed. The best-fit regression line was drawn. The Pearson’s correlation coefficient and *p-*value are provided within the panel. **Note**: ADHD – attention-deficit/hyperactivity disorder; ASD – autism spectrum disorder; BD – bipolar disorder; MDD – major depressive disorder; OCD – obsessive compulsive disorder; SCZ – schizophrenia

## Discussion

Our analysis of summary statistics from the ENIGMA ADHD, ASD, BD, MDD, OCD, SCZ and epilepsy Working Groups and the predominantly PGC case-control GWAS identified two novel findings. First, we found substantial correlations for some disorders in the pattern of sMRI case-control differences across subcortical and cortical regions. Second, these cross-disorder correlations in SBRV could partly be explained by the genetic correlations reported for these disorders from genomewide association studies (3).

The cross-disorder correlations in SBRV are intriguing because, like cross-disorder genetic correlations, they suggest that these disorders, to varying degrees, share aspects of their etiology and pathophysiology. Any interpretation of the cross-disorder sMRI correlations must keep in mind that, for all disorders, the case-control differences in sMRI measures are small (Figure 1). The largest Cohen’s d values are only −0.5 for SCZ (19, 20), −0.4 for epilepsy (14), −0.3 for BD (12, 13), −0.2 for ADHD (9, 10) and ASDs (11), and −0.1 for MDD (15, 16) and OCD (17, 18). These small case-control differences are consistent with results from GWAS and environmental risk studies, which speaks to the fact that the effects of common risk factors are, with some rare exceptions, individually small. Although it is conceivable that these small risks could accumulate to create a more dramatic pathophysiology in the brain, the ENIGMA data show that this is not the case for sMRI measures. Consistent with this finding, interindividual differences in neuroimaging account for only a small amount of the variance in symptom expression or behavioral measures of symptomatic or behavioral variance (23).

The most prominent case-control differences in cortical thickness/surface area and subcortical volumes were observed for SCZ (19, 20) and BD (12, 13). These disorders also had the highest sMRI phenotype correlations and both also showed strong sMRI phenotype correlations with MDD (15, 16) and OCD (17, 18). As Figure 2 shows, these disorders clustered together in the three-dimensional configuration required to capture cross-disorder sMRI phenotype similarity. The high sMRI correlation between SCZ and BD is consistent with prior reports of sMRI similarities between the two disorders (24). Moreover, a large body of literature reports substantial etiologic overlap between the two disorders (25-29). Because of such data, the SCZ and BD have been described as sharing a continuum of etiology leading to psychotic (30), neurophysiological (30) and neurocognitive (31) symptoms. The ENPACT study (32) showed shared fronto-temporo-occipital grey matter volume deficits in the right hemisphere of two disorders. A systematic review of associations between functional MRI activity and polygenic risk for SCZ and BD (26) reported that genetic load for these disorders affects task-related recruitment of predominantly frontal lobe brain regions.

Many studies have reported that OCD can be a comorbid diagnosis with SCZ or that patients with SCZ can have OCD symptoms (33-40). Presented findings of a significant overlap in sMRI phenotypes along with the known SCZ/OCD genetic correlations suggests that more work should examine shared pathophysiologic features between these disorders and should assess the degree to which confounds, such as medication status or chronicity, might explain these results.

The sMRI phenotype correlations mirror, to some extent, the cross-disorder correlations from genomewide association studies. Figure 3 shows a modest, yet distinct, linear correlation between the sMRI phenotype and genetic correlations. In the upper right-hand section of the plot, we see disorders having high genetic and high sMRI correlations. These are SCZ/BD, SZ/MDD, BD/MDD, OCD/BD and OCD/MDD. The inclusion of MDD in this group is notable given that it is part of the bipolar diagnosis and often occurs comorbid with other disorders. MDD also has a high genetic correlation with ADHD but a negative sMRI correlation, which makes that pair an outlier in Figure 3.

In the lower left region of Figure 3, we see disorders with low genetic and low sMRI correlations. These involve correlations of epilepsy, and correlations of ADHD with all disorders except ASDs and MDD, although the latter is somewhat of an outlier. ASDs tend to have both modest genetic correlations and modest sMRI correlations with most other disorders and, hence, populates the middle range of the figure. Like the sMRI correlations among disorders, all genetic correlations with epilepsy are low, which is consistent with the low genetic correlation between neurological and psychiatric disorders as reported by Anttila et al. (2).

The finding that SBRV correlations are correlated with genetic correlations suggests that future studies of SBRV should consider genetic sources of etiology. Yet, because only about 24% of the variance in the SBRV correlations can be accounted for by the genetic correlations, environmental sources of etiology and disease-specific genetic contributions must also be considered. These include shared confounders, such as chronicity and medication exposure, along with shared etiologic events such as birth complications or exposure to toxins *in utero*. Our prior studies of SBRV in ADHD implicated the regulation of genes in apoptosis, autophagy and neurodevelopment pathways in ADHD (41, 42). Neurodevelopmental pathways had also been implicated in the cross-disorder analysis of the Psychiatric Genomics Consortium (3), which suggests that cross-disorder similarities in these pathways may account for cross disorder similarities in SBRV.

Although we used data derived from very large samples (ENIGMA, iPSYCH and the PGC), several limitations moderate the strength of our conclusions. We inherit all the limitations of the constituent studies, but are further limited because we analyzed summary statistics, not the original data, which would require the sharing of individual subject level data, an ongoing effort among the ENIGMA disorder working groups. Thus, we cannot determine whether the possible use of controls shared among studies affected our results. It is also possible that some research participants were included in the genetic and sMRI data sets for the same disorder.

Another problem is that we could not address effects of medications or chronicity on brain structure. Furthermore, for some of the disorders, we could use youth and adult data, whereas for others only adult effect data were used. Because findings can differ substantially depending on the age range of the samples included (e.g., (9, 10, 17, 18)), this might have influenced our findings. For these reasons, analyses of participant level data will be needed to address these issues to draw stronger and more detailed conclusions. We also did not have any longitudinal data available, which limits the ability to test hypotheses about shared and unique developmental trajectories among disorders.

Despite these limitations, we have documented cross-disorder correlations in SBRV as assessed by sMRI. These cross-disorder SBRV correlations are positively associated with the disorders’ corresponding cross-disorder genetic correlations. This finding is a novel contribution worthy of further study. Our work supports conclusions from previous GWAS studies suggesting a partially shared etiology and pathophysiology among many disorders (2, 43). Disorders like SCZ and BD or ADHD and ASD, which are distinct in the diagnostic nomenclature, show significant overlap in etiology and pathophysiology. Further studies are needed to discern why brain regions are selectively affected by the risk factors that cause sMRI abnormalities (41, 42) and why these effects are correlated across disorders. Such studies may give insights into new treatment targets.

## Data availability

### URLs for GWAS

SCZ from ckqny.scz2snpres.gz (https://www.med.unc.edu/pgc/results-and-downloads)

ASD from iPSYCH-PGC_ASD_Nov2017.gz (https://www.med.unc.edu/pgc/results-and-downloads)

OCD from PGC_OCD_Aug2017-20171122T182645Z-001.zip > ocd_aug2017.gz (https://www.med.unc.edu/pgc/results-and-downloads)

ADHD from adhd_jul2017.gz (https://www.med.unc.edu/pgc/results-and-downloads)

BD from daner_PGC_BIP32b_mds7a_0416a.gz (https://www.med.unc.edu/pgc/results-and-downloads)

Epilepsy from all_epilepsy_METAL.gz (http://www.epigad.org/gwas_ilae2018_16loci.html)

MDD from PGC_UKB_depression_genome-wide.txt (http://dx.doi.org/10.7488/ds/2458)

## Acknowledgment

Dr. Faraone is supported by the European Union’s Horizon 2020 program for the CoCa project (grant agreement no 667302). and NIMH grants 5R01MH101519 and U01 MH109536-01. Research Council of Norway (#223273). Dr. Franke is supported by a personal Vici grant from the Netherlands Organization for Scientific Research (NWO, grant number 91813669) and by a grant from the European Union’s Horizon 2020 program for the CoCa project (grant agreement no 667302). ENIGMA work is supported by NIH grants U54 EB020403 (PI: Thompson), R01 MH116147 (PI: Thompson) and R01MH117601 (MPIs: Jahanshad & Schmaal). Dr Hoogman is supported by a personal Veni grant from the Netherlands Organization for Scientific Research (NWO, grant number 91619115). Dr. Mcdonald is supported by NIH grants R01 NS065838 and R21 NS107739. P.Rovira is a recipient of a pre-doctoral fellowship from the Agència de Gestió d’Ajuts Universitaris i de Recerca (AGAUR), Generalitat de Catalunya, Spain (2016FI_B 00899). Dr. Schmaal is supported by a NHMRC Career Development Fellowship (1140764). Dr Sisodiya is supported by Epilepsy Society, UK, and the work was partly undertaken at UCLH/UCL, which received a proportion of funding from the UK Department of Health’s NIHR Biomedical Research Centres funding scheme.). Dr. Van Erp is supported by NIH grants U54 EB020403 (PI: Thompson), R01 MH116147 (PI: Thompson), R01MH117601 (MPIs: Jahanshad & Schmaal), and R01MH121246 (MPIs: Turner, Van Erp, & Calhoun).

