## Supplementary file for "Structural Brain Imaging Studies Offer Clues about the Effects of the Shared Genetic Etiology among Neuropsychiatric Disorders"

**Supplemental Figure 1: Leave One Disorder Out Sensitivity Analysis**

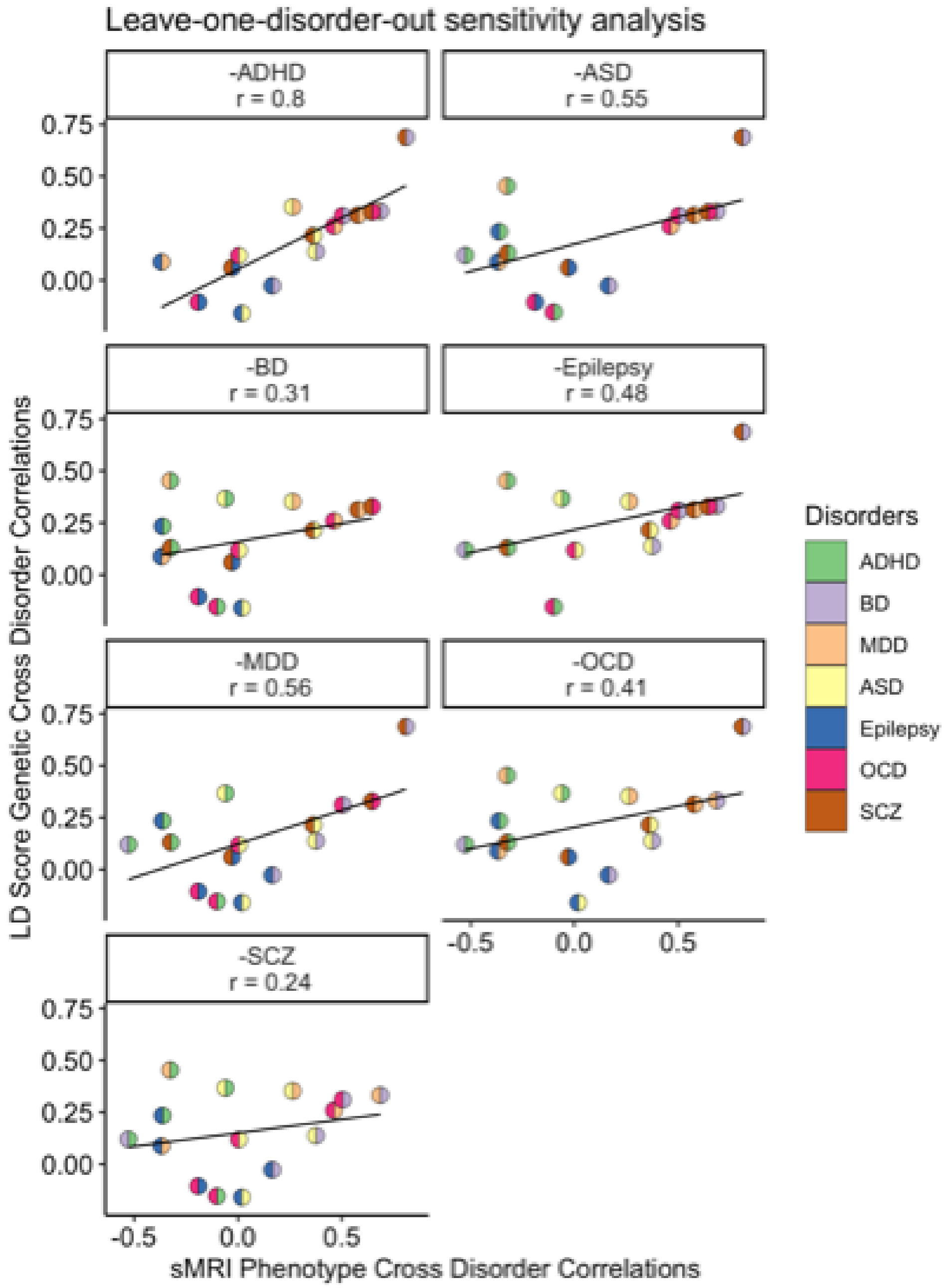

**Figure legend**: Scatter plot showing sensitivity analyses for the correlation of correlations analyses. Genetic correlations ($r_g$) computed by LD-score regression are on the vertical axis, with correlations of Cohen's d values displayed on the horizontal axis. Each dot is color-coded according to the pair-wise disorder correlations that were computed. The best-fit regression line was drawn. The Pearson's correlation coefficient and *p*-value are provided within the panel.

**Note**: ADHD – attention-deficit/hyperactivity disorder; ASD – autism spectrum disorder; BD – bipolar disorder; MDD – major depressive disorder; OCD – obsessive compulsive disorder; SCZ – schizophrenia.

**Supplementary Table 1.** Covariate Adjustments Used by Each Study

| **Disorder** | **Brain measure** | **Covariates** |
|---|---|---|
| Bipolar disorder | Cortical thickness and surface area | Age, sex, scan center, ICV (surface area only) |
| Bipolar disorder | Subcortical volume | Age, sex, ICV |
| Depression | Cortical thickness and surface area | Age, sex, scan center |
| Depression | Subcortical | Age, sex, scan center, ICV |
| Schizophrenia | Cortical thickness and surface area | Age, sex |
| Schizophrenia | Subcortical | Age, sex, ICV |
| OCD | Cortical thickness and surface area | Age, sex, scan center, ICV (surface area only) |

| | | |
|---|---|---|
| OCD | Subcortical | Age, sex, scan center, ICV |
| ASD | Cortical thickness and surface area | Age, sex, scan center<br><br>(groups matched on ICV) |
| ASD | Subcortical | Age, sex, scan center<br>(groups matched on ICV) |
| Epilepsy | Cortical thickness and surface area | Age, sex, ICV |
| Epilepsy | Subcortical | Age, sex, ICV |
| ADHD | Cortical thickness and surface area | Age, sex, scan center, ICV (surface area only) |
| ADHD | Subcortical | Age, sex, ICV |

**Supplementary Table 2.** Additional information pertaining to the genomewide association studies (GWASs)

| Disorder | Number of cases | Number of controls | Estimated SNP-$h^2$ (SE) | References | Link to dataset |
|---|---|---|---|---|---|
| ADHD | 20,183 | 35,191 | 0.23 (0.014) | Demontis et al., 2019 (Nat Genet) | https://www.med.unc.edu/pgc/results-and-downloads |
| Autism spectrum disorder | 18,382 | 27,969 | 0.20 (0.016) | Grove et al., 2019 (Nat Genet) | https://www.med.unc.edu/pgc/results-and-downloads |
| Bipolar disorder | 20,352 | 31,358 | 0.35 (0.016) | Stahl et al., 2019 (Nat Genet) | https://www.med.unc.edu/pgc/results-and-downloads |
| Epilepsy | 15,212 | 29,677 | 0.11 (0.017) | The International League Against Epilepsy Consortium on Complex Epilepsies, 2018 (Nat Commun | http://www.epigad.org/gwas_ilae2018_16loci.html |
| Major depressive disorder | 170,756 | 329,443 8 | 0.06 (0.0023) | Howard et al., 2019 (Nat Neurosci) | http://dx.doi.org/10.7488/ds/2458 (MDD dataset excluding 23andMe) |
| Obsessive compulsive disorder | 2,688 | 7,037 | 0.33 (0.048) | International Obsessive Compulsive Disorder Foundation Genetics Collaborative (IOCDF-GC) and OCD Collaborative Genetics Association Studies (OCGAS) 2018 (Mol Psychiatry) | https://www.med.unc.edu/pgc/results-and-downloads |
| Schizophrenia | 36,989 | 113,075 | 0.24 (0.009) | Ripke et al., 2014 (Nature) | https://www.med.unc.edu/pgc/results-and-downloads |
